# MolPad: An R-Shiny Package for Cluster Co-Expression Analysis in Longitudinal Microbiomics

**DOI:** 10.1101/2023.11.29.569321

**Authors:** Kaiyan Ma, Margaret W. Thairu, Kris Sankaran

## Abstract

The R-Shiny package MolPad provides an interactive dashboard for understanding the dynamics of longitudinal molecular co-expression in microbiomics. The main idea for addressing the issue is first to use a network to overview major patterns among their predictive relationships and then zoom into specific clusters of interest. It is designed with a focus-plus-context analysis strategy and automatically generates links to online curated annotations. The dashboard consists of a cluster-level network, a bar plot of taxonomic composition, a line plot of data modalities, and a table for each pathway. Further, the package includes functions that handle the data processing for creating the dashboard. This makes it beginner-friendly for users with less R programming experience. We illustrate these methods with a case study on a longitudinal metagenomics analysis of the cheese microbiome.

## Statement of need

The realm of microbiomics is expanding rapidly, with numerous new studies and methodologies emerging (Bokulich et al. 2020). This highlights the need for visual exploration tools that can account for interaction across biological modalities (Fernstad et al. 2011). It’s important to enable interpretations of dynamics and network structure because these have specific meanings in the genomic context (Corel et al. 2016). Another issue is the annotation of notable features. A characteristic of microbiome data is that each identical feature can be classified at several levels of taxonomic resolution and could have several IDs in different databases (Kanehisa and Sato 2020). Although relevant annotation is typically available online, it can be tedious to search through databases manually. Moreover, microbiome data often exhibit longitudinal variation. In this context, we must gain insight into the functioning of how individual features change and how they may influence related features. These issues have posed a challenge for unified visualization and interpretation.

In response to the above issues, previous studies on interactive visualization tools have designed methods to work on such data. `microViz` (Barnett, Arts, and Penders 2021) provides a Shiny app for interactive exploration by pairing ordination plots and composition circular bar charts to show each taxon’s prevalence and abundance. `GWENA` (Lemoine et al. 2021) applies a network in conducting gene co-expression analysis and extended module characterization in a single package to understand the underlying processes contributing to a disease or a phenotype. `NeVOmics` (Zúñiga-León, Carrasco-Navarro, and Fierro 2018) improved compatibility with a dynamic dashboard and facilitated the functional characterization of data from -omics technologies. It also integrates over-representation analysis and network-based visualization to display enrichment results. We build on this foundation to better enable longitudinal and cluster-oriented visualization.

## Methods

### Network Generation

We first scale and cluster the trajectories across all molecular features to depict longitudinal change. For clustering, we use K-means and a built-in elbow method to choose the optimal number. Then, we predict a co-expression network for the extracted patterns, similar to how GENIE3 (Huynh-Thu 2010) creates gene regulatory networks. We also divide the prediction process into individual regression tasks. Each cluster centroid is predicted from the expression patterns of all the other cluster centroids, using random forests. We choose random forests because of their potential to model interacting features and non-linearity without strong assumptions. The Mean Decrease Accuracy of a subset of top predictors whose expression directly influences the expression of the target cluster is taken as an indication of a putative link. That is to say, based on the random forest prediction, if two groups of features are highly linked according to the network, they will have strongly related longitudinal patterns, as shown in Fig 3.

### Network Navigation

Navigating the network in the MolPad dashboard follows three steps, as shown in Fig 1: First, choose a primary functional annotation. Adjustment options for fine-tuning include network layout and importance threshold for edge density. Bright green nodes (Fig 2.A) represent clusters containing the most features in the chosen functional annotation. Second, brushing on the network reveals patterns of taxonomic composition (Fig 2.B) and typical trajectories (Fig 2.C). The user can also zoom into specific taxonomic annotations by filtering. Third, they may view the feature table (Fig 2.D), examine the drop-down options for other related function annotations, and click the link for online details for the items of interest. The interface is designed to support iterative exploration, encouraging the use of several steps to answer specific questions, like comparing the distributional patterns between two functions or finding functionally important community members metabolizing a feature of interest. By applying a focus-plus-context approach ((Shneiderman 1996) and (Sankaran and Holmes 2018)), we can bridge the examination of high-level details related to individual features with contextual information about cluster interactions within the network visualization.

**Figure 1.**
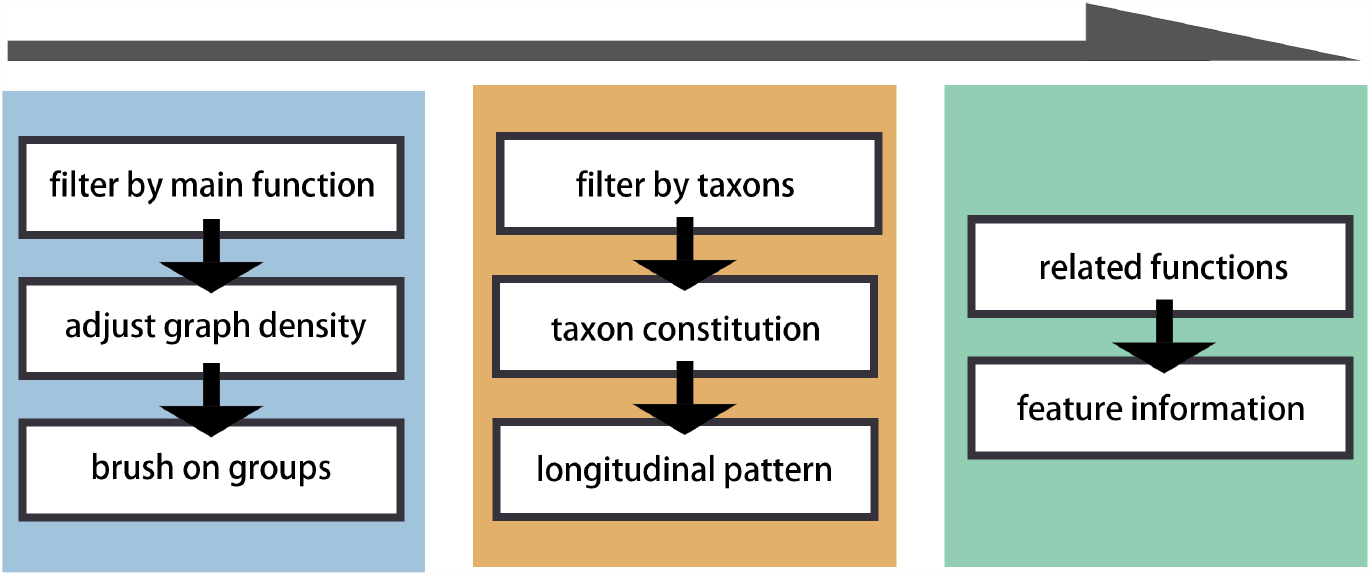
Overview and workflow of using MolPad package.

**Figure 2.**
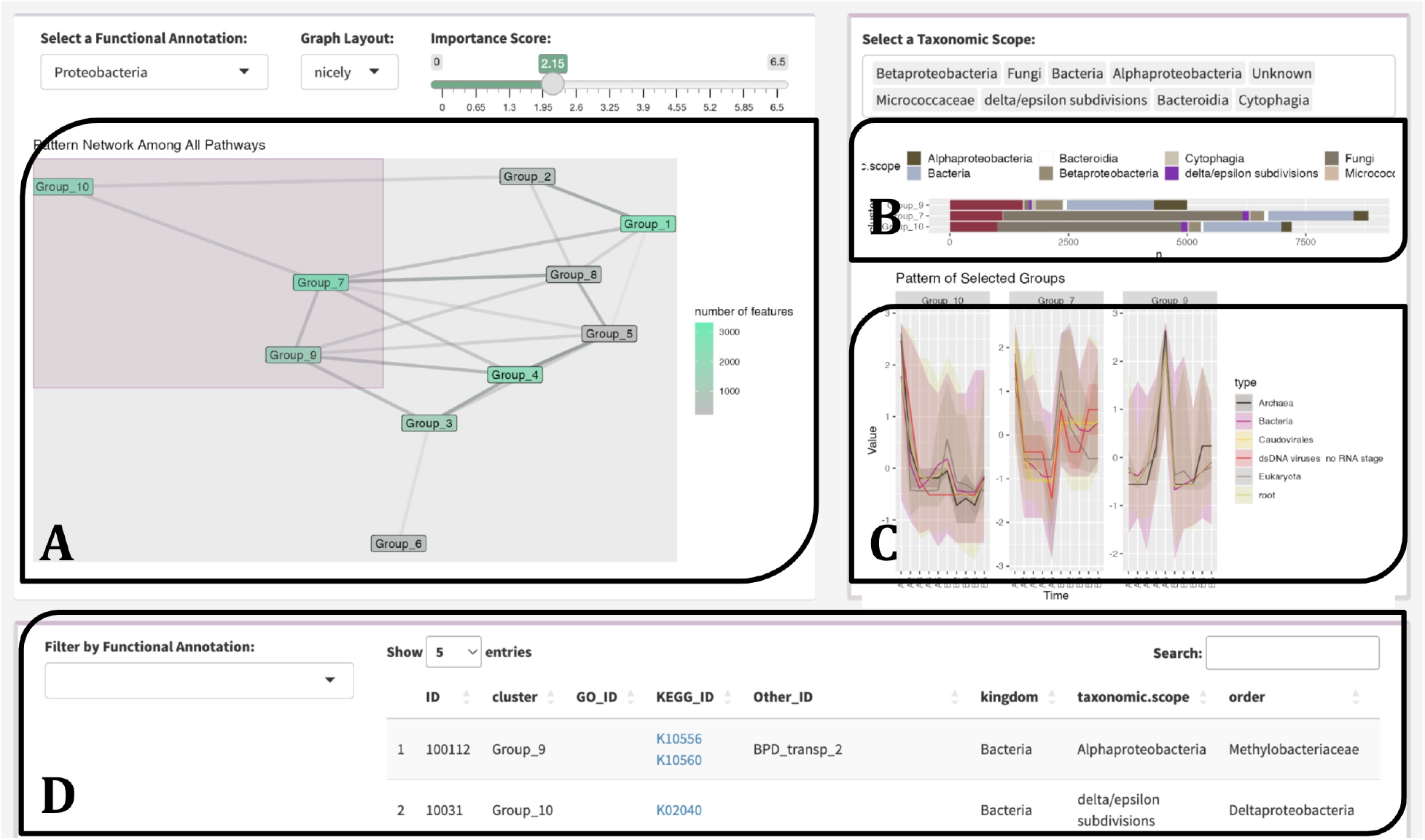
Dashboard Overview: `A`: cluster-level network, `B`: taxonomic-level bar plot, `C`: a type-level line plot, and `D`: a feature-level table.

**Figure 3.**
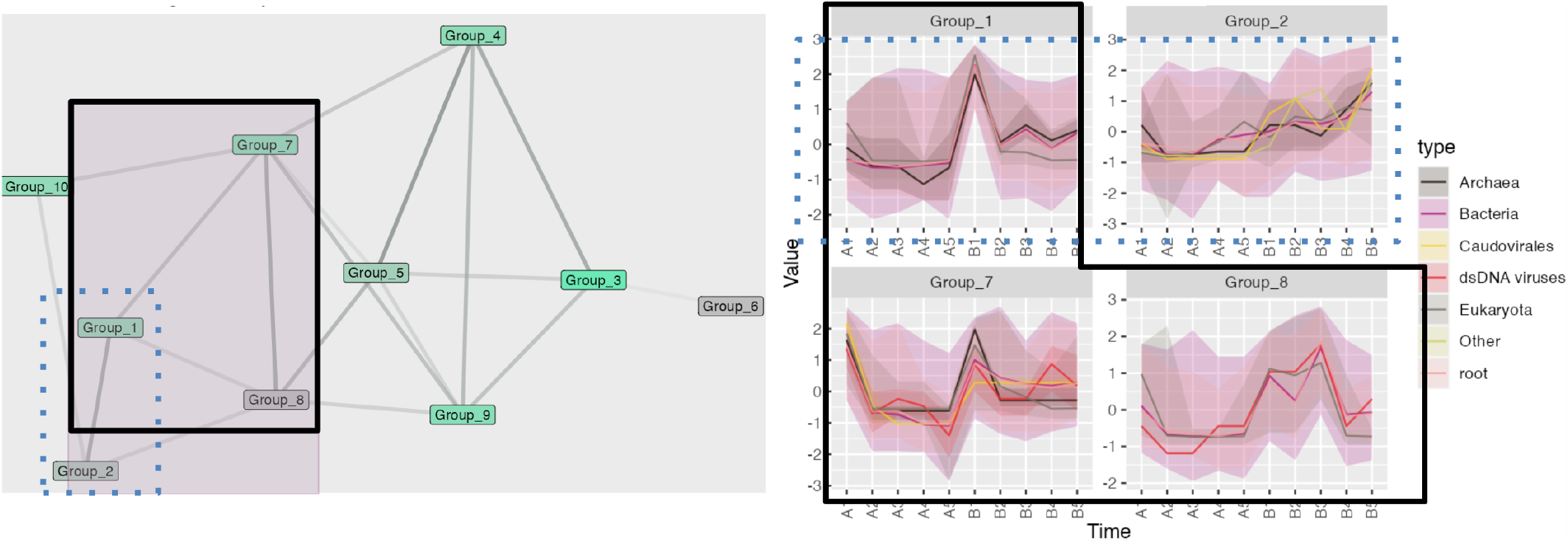
Example of discovering related patterns with the network plot. The shade of edges stands for the vicinity of nodes. In the brushed area, Groups 1-7-8 (circled by solid black lines) and 1-2 (circled by blue dashed lines) are strongly linked. For Groups 1, 7, and 8, the patterns are w-shape with an evident peak at the same time section. For Groups 1 and 2, although Group 1 has higher volatility, they both show a highly overlapped increasing trend.

**Figure 4.**
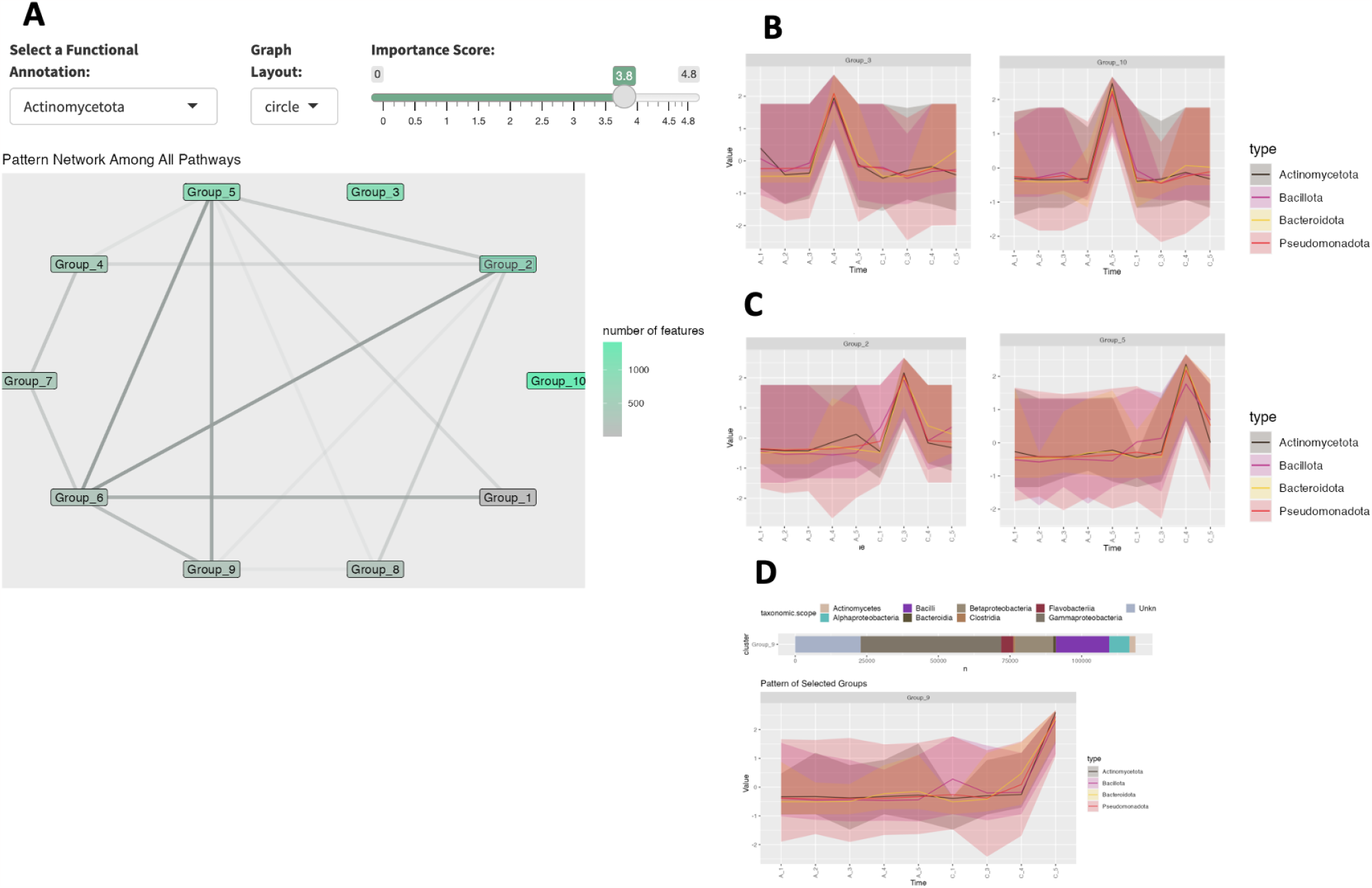
Dashboard showing Actinomycetota filtered network (A) with enrichment pattern for Cheese Sample-A (B) and Cheese Sample-C (C); Cluster pattern for Group 9, which also is enriched for Type IV secretion genes (D).

### Case Study: Cheese Data

Here we aim to highlight the versatility of the MolPad Dashboard with a case study of microbial communities on the wash-rind cheese’ surface collected during cheese ripening. In the original research, (Saak et al. 2023) investigated the successional dynamics that occur within cheese rind microbial communities using a combination of 16S rRNA amplicon, Illumina, and PacBio sequencing. We functionally and taxonomically annotate (using eggNOG (Huerta-Cepas et al. 2018) and MMseqs2 (Steinegger and Söding 2017)) the contigs they have generated from the Illumina reads, to demonstrate the utility of MolPad. Specifically, we focus on Cheese Sample A and Cheese Sample C. Overall, we are not only able to uncover similar findings as (Saak et al. 2023), with this subset, but we are also able to make new hypotheses.

For example, when we filter for Actinomycetota (Actinobacteria) as the functional group, we see that there are no edges connecting to group 10 and group 3-the clusters that have the most features associated with Actinomycetoa for Cheese sample A (4.A). Looking at the pattern traces of these groups, (4.B), there is a peak in samples A4 (week 9) and A5 (week 13), which mirrors the 16S rRNA results of Saak et al. Since these two clusters do not have edges connecting them to other groups, this suggests that the Actinomycetoa features found in these groups follow distinct longitudinal succession patterns that are independent. When looking at Actinomycetoa within Cheese Sample C we see a different pattern. Groups 2 and 5, have the most features associated with Actinomycetoa, but they are highly connected to the other groups (4.A). From these results, we can hypothesize that though Actinomycetoa features are more abundant in later time points for both cheese samples, their dynamics are differentially influenced. The authors found that Type VI secretion was enriched in Pseudomonadota bacteria (specifically, *Psychrobacter*), and hypothesized this enrichment was due to the importance of physical species interactions that occur with this habitat. Using MolPad, we searched for other secretion systems associated genes, to understand their dynamics within the community. Focusing on KEGG annotated Type IV secretion genes, we found that Group 9 contained 12/13 of these genes. Within this group, features that cluster are ones that peak in Cheese sample C5 (week 13, 4.D). This is also the most taxonomically diverse sample. From this, we can hypothesize that increased taxonomic diversity is also associated with increases in genes that are related to competitive species interactions.

### Usage

The source code for `MolPad` is available on Github. The app is hosted in the R package which can be downloaded and run locally. We anticipate that some users may need more flexibility in their analyses, requiring backend R development for tasks like setting up detailed operating models or downloading figure outputs. For such needs, the essential set of R functions employed in the Shiny app is accessible through the R package.

## Supporting information

Github Link for Supplemental-Materials

## Acknowledgments

Research reported in this publication was supported by the National Institute of General Medical Sciences of the National Institutes of Health under award number R01GM152744.

## Notes

https://github.com/KaiyanM/MolPad

### Competing Interest Statement

The authors have declared no competing interest.

